# Positively selected effector genes and their contribution to virulence in the smut fungus *Sporisorium reilianum*

**DOI:** 10.1101/177022

**Authors:** Gabriel Schweizer, Karin Münch, Gertrud Mannhaupt, Jan Schirawski, Regine Kahmann, Julien Y. Dutheil

## Abstract

Plants and fungi display a broad range of interactions in natural and agricultural ecosystems ranging from symbiosis to parasitism. These ecological interactions result in coevolution between genes belonging to different partners. A well-understood example are secreted fungal effector proteins and their host targets, which play an important role in pathogenic interactions. Biotrophic smut fungi (Basidiomycota) are well-suited to investigate the evolution of plant pathogens, because several reference genomes and genetic tools are available for these species. Here, we used the genomes of *Sporisorium reilianum* f. sp. *zeae* and *S. reilianum* f. sp. *reilianum*, two closely related formae speciales infecting maize and sorghum, respectively, together with the genomes of *Ustilago hordei*, *Ustilago maydis* and *Sporisorium scitamineum* to identify and characterize genes displaying signatures of positive selection. We identified 154 gene families having undergone positive selection during species divergence in at least one lineage, among which 77% were identified in the two investigated formae speciales of *S. reilianum*. Remarkably, only 29% of positively selected genes encode predicted secreted proteins. We assessed the contribution to virulence of nine of these candidate effector genes in *S. reilianum* f. sp. *zeae* by deleting individual genes, including a homologue of the effector gene *pit2* previously characterized in *U. maydis*. Only the *pit2* deletion mutant was found to be strongly reduced in virulence. Additional experiments are required to understand the molecular mechanisms underlying the selection forces acting on the other candidate effector genes, as well as the large fraction of positively selected genes encoding predicted cytoplasmic proteins.

## Introduction

Plants and fungi have a long history of coevolution since the emergence of pioneering land plants approximately 400 million years ago. The development of early plants was likely supported by associations with symbiotic fungi, as suggested by analyses of ribosomal RNAs and fossil records (Remy et al. 1994; Gehrig et al. 1996; Martin et al. 2017). Different forms of plant-fungus interactions have evolved, including mutualistic symbiosis where both plant and fungus benefit (Parniske 2008), and pathogenic interactions where fungal colonization greatly reduces plant fitness (Dean et al. 2012). Pathogenic interactions play critical roles in natural and agricultural ecosystems, and understanding the evolutionary mechanisms shaping them is of great importance to plant production, food security and protection of biodiversity in natural ecosystems (Fisher et al. 2012; Bagchi et al. 2014).

Secreted fungal effector proteins are key players in pathogenic interactions as they are involved in protecting and shielding growing hyphae, suppressing plant defense responses and changing plant physiology to support growth of the pathogen (Stergiopoulos and de Wit 2009; de Jonge et al. 2011; Giraldo and Valent 2013). Many effector proteins lack known functional domains, and expression of a subset of effectors is linked to plant colonization (Lo Presti et al. 2015; Toruno et al. 2016; Franceschetti et al. 2017; Lanver et al. 2017). Effector proteins with a strong effect on virulence phenotype are thought to coevolve with their plant targets either in an arms race or a trench-warfare scenario (Brown and Tellier 2011). In the former, fungal effectors manipulating the host are under positive directional selection, and plant targets evolve in response to changes in effector proteins (Rovenich et al. 2014). In the latter scenario, sets of alleles are maintained by balancing selection in both host and pathogen populations (Brown and Tellier 2011; Tellier et al. 2014). Several methods are available for identifying genomic regions under selection (Nielsen 2005; Aguileta et al. 2009; Aguileta et al. 2010). It has been proposed that genes with signatures of positive selection have important functions during host pathogen interaction or have contributed to host specialization (Tiffin and Moeller, 2006). It is therefore expected that the deletion of such genes reduces virulence when tested on a susceptible host.

Depending on the aim of the investigation, studies identifying genes with signatures of positive selection are carried out within or between species. Whilst studies on the population level focus on recent and ongoing selective processes and are instrumental in the understanding of adaptation, comparative genomic studies employing different species encompass a broader time span and provide insight into the underlying genetic basis of host specialization (Plissonneau et al. 2017). The signature of positive selection in such case typically takes the form of an excess of divergence between species due to increased fixation of mutations by selective sweeps compared to a neutral expectation (Yang and Nielsen, 1998). This is commonly measured by the ratio of non-synonymous (d_N_) over synonymous (d_S_) divergence and a d_N_ / d_S_ ratio > 1 is taken as evidence for positive selection under the assumption that synonymous substitutions are neutral while non-synonymous are not. Positive selection studies in a number of plant pathogen systems revealed that genes encoding secreted effector proteins are enriched in signatures of positive selection (Möller and Stukenbrock 2017). Such studies include investigations in diverse plant pathogens like *Microbotryum* species causing anther-smut disease of Caryophyllaceae species (Aguileta et al. 2010), the wheat pathogen *Zymoseptoria tritici* (Stukenbrock et al. 2011), the rust fungus *Melampsora larici-populina* (Hacquard et al. 2012), the rice blast fungus *Magnaporthe oryzae* (Huang et al. 2014), the wheat stem rust fungus *Puccinia graminis* f. sp. *tritici* (Sperschneider et al. 2014) the Irish potato famine pathogen *Phytophthora infestans* (Dong et al. 2014), a group of *Fusarium* species (Sperschneider et al. 2015) and a group of smut fungi parasitizing different grasses and a dicot host (Sharma et al. 2014; Sharma et al. 2015). Yet, as only a few genes under positive selection have been functionally studied, the link between the selected genotypes and their corresponding phenotypes are only beginning to be understood and only a few studies used evolutionary predictions to unravel the molecular mechanisms of host adaptation. For example, the population genomics study in the wheat pathogen *Z. tritici* which identified candidate effector genes under positive selection (Stukenbrock et al. 2011) was followed up experimentally, and in this case it was shown that the deletion of three of four candidate genes reduced virulence (Poppe et al. 2015). In the grey mold fungus *Botrytis cinerea* four positively selected genes were deleted without affecting virulence, and this finding was attributed to functional redundancy, the limited number of tested host plants, or experimental conditions different from natural infections (Aguileta et al. 2012). A study of the oomycete effector protein EpiC1 showed that a single amino acid substitution at a site under positive selection affected the binding affinity of different host proteases determining host specificity (Dong et al. 2014).

Smut fungi, belonging to the division of Basidiomycota, are a group of about 550 species parasitizing mostly grasses, including important crops like maize, sorghum, oat, barley and sugarcane (Begerow et al. 2014). In smut fungi, sexual reproduction is linked to pathogenic development and smut fungi therefore depend on successful plant colonization to complete their life cycle. As biotrophic pathogens, they require living plant tissue for establishing a successful interaction (Martinez-Espinoza et al. 2002). With few exceptions like *Ustilago maydis*, smut fungi usually develop symptoms only in the female or male inflorescence of their respective host plants. During the last ten years, quality draft genome sequences of prominent species were obtained, including *U. maydis*, the causative agent of smut disease on maize and teosinte (Kämper et al. 2006), *Sporisorium reilianum* causing head smut of maize and sorghum (Schirawski et al. 2010), *Ustilago hordei* infecting barley (Laurie et al. 2012), and *Sporisorium scitamineum* parasitizing sugarcane (Que et al. 2014; Taniguti et al. 2015;Dutheil et al. 2016). The head smut fungus *S. reilianum* occurs in two formae speciales that infect maize (*S. reilianum* f. sp. *zeae*) or sorghum (*S. reilianum* f. sp. *reilianum*) (Zuther et al. 2012). The concept of formae speciales is used in phytopathology to distinguish members of the same species based on their ability to colonize a certain host plant (in this example maize or sorghum) (Anikster, 1984). The divergence of *U. hordei*, *U. maydis*, *S. scitamineum* and *S. reilianum* was inferred to have occurred in the interval of seven and 50 million years ago (Munkacsi et al. 2007). The availability of genome sequences of several species with different host ranges, together with established tools for genetic manipulations (Brachmann et al. 2004; Kämper 2004; Khrunyk et al. 2010; Schuster et al. 2016) make this group of smut fungi particularly interesting to study the evolution of effector genes as well as their contributions to virulence, speciation and host specificity.

Here, we employed the genome of the recently sequenced strain *S. reilianum* f. sp. *reilianum* SRS1_H2-8 (http://www.ebi.ac.uk/ena/data/view/LT795054-LT795076) (Zuther et al. 2012) together with the genomes of *U. maydis*, *U. hordei*, *S. scitamineum* and *S. reilianum* f. sp. *zeae* to identify potential effector genes with signatures of positive selection. Candidate genes were individually deleted in *S. reilianum* f. sp. *zeae* and the phenotype of the deletion strains was assessed after infection of maize in order to understand their function with respect to virulence. We report that the deletion of one candidate gene, *pit2*, led to a strong reduction in virulence and we further discuss hypotheses on the origin of positive selection for the other candidate genes.

## Materials and Methods

### Construction of homologous protein families

Fungal species used in this study, their number of gene models, number of predicted secreted proteins and sources of genome data are listed in supplementary table S1. The predicted proteome of the five smut fungi *U. hordei*, *U maydis*, *S. scitamineum*, *S. reilianum* f. sp. *zeae* and *S. reilianum* f. sp. *reilianum* were used to perform an all-against-all blastp search (Altschul et al. 1990). The SiLiX algorithm was subsequently used to infer homology relationships based on the blast hits (Miele et al. 2011). Two parameters are considered to decide whether a Blast hit can be taken as evidence for homology: the percent identity between two sequences and the relative length of the hit compared to the total length of the two sequences, hereby referred to as “coverage”. In order to maximize the number of families comprising 1:1 orthologues (that is families that have an equal number of members in each species), SiLiX (Miele et al. 2011) was run with a range for coverage and identity thresholds between 5 % and 95 % in 5 % steps. An identity of 40 % and coverage between 5 % and 45 % lead to the maximum number of families with 1:1 orthologues (5,394; supplementary fig. S1) while settings with 40 % identity and 80 % coverage lead to 5,326 families with 1:1 orthologues (supplementary fig. S1). Since using a higher coverage had only a cost of 68 core families, the stricter criteria were applied for family clustering. Families with at least two members were aligned on the codon level using MACSE 1.01b (Ranwez et al. 2011) and on the protein level using PRANK v.100802 (Löytynoja and Goldman 2008). The resulting alignments were subsequently compared and column scores (CS) computed for each position in the alignment (Thompson et al. 1999). Only positions with CS of 100 % (that is, alignment columns identically found by both methods) and a maximum of 30 % gaps were retained for further analysis.

### Estimation of genome-wide divergence values and divergence times

The five genomes of *U. hordei*, *U. maydis*, *S. scitamineum*, *S. reilianum* f. sp. *zeae* and *S. reilianum* f. sp. *reilianum* were aligned using the Multiz genome aligner from the TBA package (Blanchette et al. 2004) and projected on the *U. maydis* genome as reference. The resulting multiple genome alignment had a total size of 21 Mb and was further restricted to regions with homologous sequences in the five species (total length after this step: 14.3 Mb) and processed to remove coding regions. The final non-coding sequence alignment had a total length of 2.2 Mb, for which pairwise nucleotide similarities were computed in non-overlapping windows of 10 kb.

Gene families with exactly one member in each species were concatenated and pairwise protein sequence similarities computed using the *seqinr* package for R (Charif and Lobry 2007). Protein alignments were also used to infer dates of divergence, using a relaxed clock model. The PhyloBayes version 4.1 software (Lartillot et al. 2009) was used with the auto-correlated model of Thorne et al. (Thorne et al. 1998) under a GTR + CAT model. A unique calibration point was used, based on the divergence time of the most divergent lineage *U. hordei*, previously estimated to have occurred between 27 and 21 Myr (Bakkeren and Kronstad, 2007). A uniform prior was used on this interval for the Monte-Carlo Markov Chain. As convergence issues arise when large alignments (more than 20,000 positions) are used, we followed the PhyloBayes authors’ recommendation to conduct a jackknife procedure. We generated three datasets of ca 20,000 amino acids by randomly sampling families and concatenating the corresponding alignments. Two chains were run in each case and convergence was assessed. Sampling was performed after a burning of 10,000 iterations, and every 10 subsequent iterations. Chains were run to ensure that the minimum effective sample size was greater than 50 and maximum relative difference lower than 0.3 in at least one sample. Results are summarized in supplementary table S2 and supplementary fig. S2 shows the six chains for the three samples. In addition to the convergence of the two chains for each sample, our results reveal extremely consistent results between samples. Figure 1A shows estimates from one chain of the third data set, which shows a minimum effective sample size greater than 300.

**FIG. 1.**
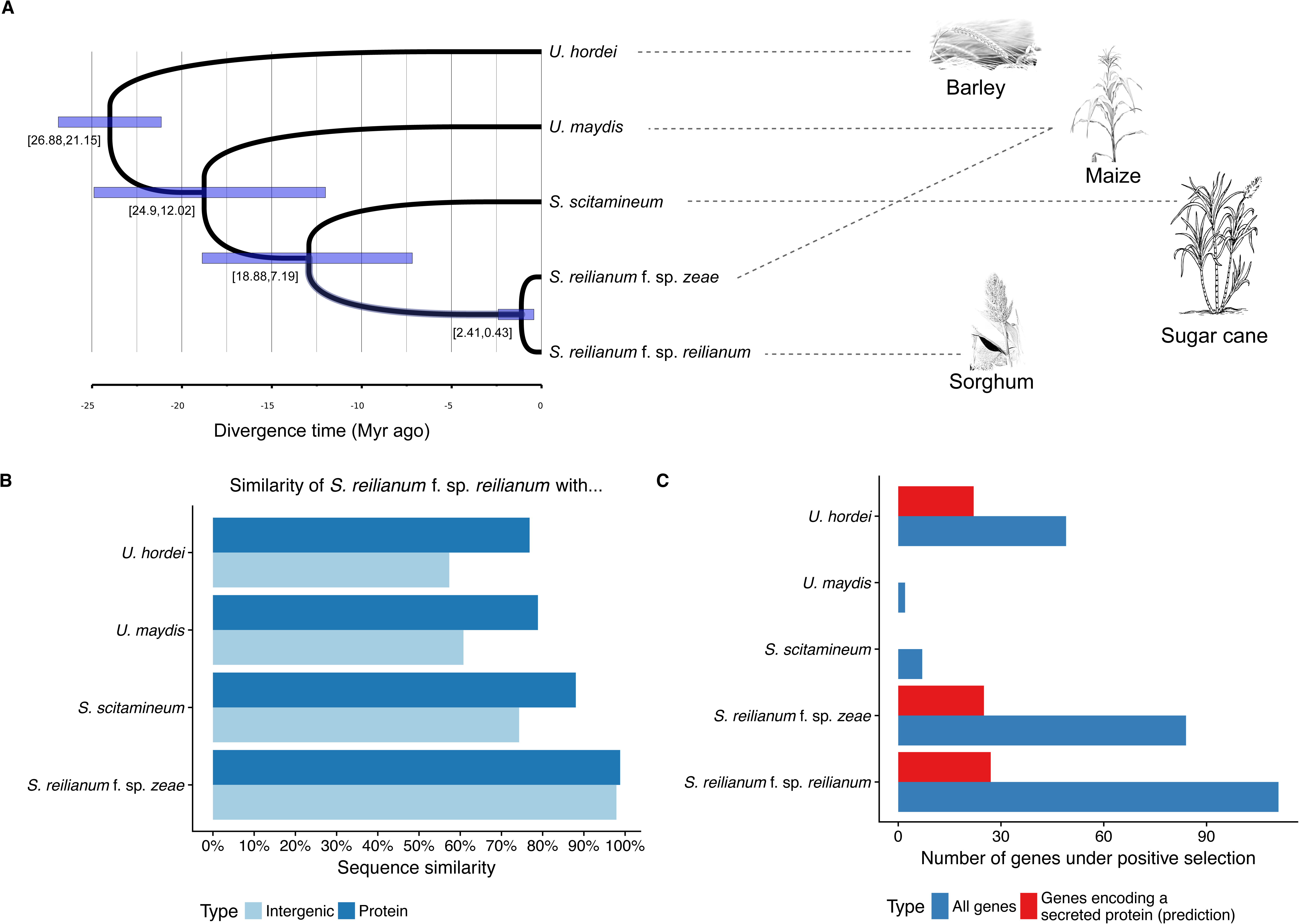
Phylogeny, divergence estimates and number of genes under positive selection in five related smut fungal species parasitizing different host plants. A) Chronogram of the five fungal pathogens as estimated under a relaxed molecular clock. Boxes represent 95% posterior intervals, with corresponding values indicated below. B) Pairwise sequence differences, for both the non-coding genome and the proteome (non-synonymous differences). C) Number of positively selected genes on each terminal branch (total number of genes and genes predicted to encode a secreted protein).

### Detection of positive selection

For gene families with at least three members, translated sequences were employed to create maximum likelihood phylogenetic trees using PhyML 3.0 (Guindon et al. 2010) with a minimum parsimony starting tree and the LG amino acid substitution model with a four-classes gamma distribution of site-specific substitution rate (Le and Gascuel 2008). The best tree topology obtained from nearest neighbor interchange (NNI) and subtree pruning recrafting (SPR) searches was kept (Guindon et al. 2010). BppML (Dutheil and Boussau 2008) was then used to re-estimate branch lengths from the codon alignment using the YN98 substitution model (Nielsen and Yang 1998). We next aimed at inferring the occurrence of positive selection for each gene family. This is typically achieved by measuring the ratio of non-synonymous vs. synonymous substitutions (d_N_/d_S_ ratio) using models of codon sequence evolution (Yang, 2006). In particular, non-homogeneous models of sequence evolution estimate the d_N_/d_S_ ratio independently in different lineages, yet at the cost of potential over-parametrization issues. In the manual of the PAML package, the authors state that such models should only be used for hypothesis testing and advise against using them for scans of positive selection. Dutheil et al. (2012) proposed a model selection approach (implemented in the TestNH package) allowing to select for the best non-homogeneous model supported by the data. They start by fitting the simplest (homogeneous) model and sequentially add parameters to model variation of selective regime among lineages. Because the number of possible models is large even for small data sets, two heuristic approaches have been introduced: the ‘free’ heuristic permits unconnected branches from the tree to evolve under the same regime, while the ‘joint’ heuristic restricts model sharing to connected branches (see Dutheil et al. 2012 for details). The choice of models to test is guided by statistics on the patterns of substitutions on the phylogenetic tree, an approach named *substitution mapping* (Romiguier et al. 2012). Apart from the model selection approach, the underlying models of codon sequence evolution are identical to the one originally described by Yang (Yang, 1998; Yang and Nielsen, 1998). Model selection was performed with the TestNH software, which contains two programs: (1) MapNH (Romiguier et al. 2012) was used for mapping substitutions on the previously inferred phylogenetic tree and (2) PartNH (Dutheil et al. 2012) was subsequently employed to fit time non-homogeneous models of codon substitutions. PartNH uses the previously inferred substitution maps in order to perform model comparisons and select a non-homogeneous model with minimal number of parameters. Both methods ‘free’ and ‘join’ were applied and compared to scan for positive selection. Finally, putative secreted effector proteins were identified by predicting secretion using SignalP 4.0 (Petersen et al. 2011) and proteins were considered as secreted if the program indicated the presence of a signal peptide but no transmembrane domain.

To detect residues under positive selection in homologues of *pit2*, the branch-site model with Bayes Empirical Bayes (BEB) analysis as implemented in PAML4 (Yang 2007) was applied. We employed information about family composition, alignment and phylogeny as outlined above and defined *sr10529* and *srs_10529* as foreground branches. A posterior probability threshold of > 95 % was used for the BEB analysis.

### Association of positively selected genes with repeats in U. hordei

We tested whether genes under positive selection are located significantly closer to repetitive elements than average genes in the genome of *U. hordei*, which shows the highest content of repetitive elements in the group of smut fungi investigated here. For this analysis, only a group of “uncharacterized interspersed repeats” was investigated, because it was shown previously that this is the only category showing a strong association with candidate effector genes (Dutheil et al. 2016). Binary logistic regressions were conducted in R using the *rms* package (Harrell 2015). The ‘robcov’ function of the rms package was used in order to get robust estimates of each effect. The variable “distance to the closest interspersed repeat” was transformed by log(x+1) because of its extreme distribution.

### Comparing d_N_/d_S_ ratios of genes residing in virulence clusters

Previous work has identified several virulence gene clusters in *U. maydis* and some of them play important roles during pathogenic development (Kämper et al. 2006; Schirawski et al. 2010). In total, these clusters contain 163 genes, where 100 reside in clusters without virulence phenotype and 63 reside in clusters with virulence phenotype upon deletion. Both types of clusters contain each 32 genes for which a d_N_/d_S_ ratio could be determined (the missing genes are part of families that do not have at least three members and were therefore not analyzed). The d_N_/d_S_ ratios of all genes in clusters were compared between clusters with and without virulence phenotype (Wilcoxon Rank-Sum Test).

### Gene Ontology terms enrichment analysis

All proteins in *S. reilianum* f. sp. *zeae*, in *S. reilianum* f. sp. *reilianum* and in *U. hordei* were considered for Gene Ontology (GO) term enrichment analyses. GO terms were assigned using iprscan 1.1.0 (http://fgblab.org/runiprscan; developed by Michael R. Thon) which links GO information provided by Interpro to each protein. In this way, 1,759 unique GO terms could be assigned to 4,130 proteins in *S. reilianum* f. sp. *zeae*, 1,744 unique GO terms could be assigned to 4,124 proteins in *S. reilianum* f. sp. *reilianum* and 1,757 unique GO terms could be assigned to 3,922 proteins in *U. hordei* (supplementary table S3). The Bioconductor package topGO (Alexa et al. 2006) was then used to link each GO term to the three major categories “Cellular Component”, “Biological Process” or “Molecular Function”. Enrichment analysis was performed by computing *P* values for each GO term using Fisher’s classic test with parent-child correction (Grossmann et al. 2007). Cytoplasmic proteins with and without signatures of positive selection were compared for the three species separately, and differences were considered to be significant at the 5 % level.

### Strains and growth conditions

The *Escherichia coli* derivative Top 10 (Invitrogen, Karlsruhe, Germany) and the *Saccharomyces cerevisiae* strain BY4741 (MATa *his*3Δ1 *leu*2Δ *met*15Δ *ura*3Δ; Euroscarf, Frankfurt, Germany; kindly provided by M. Bölker, Marburg) were used for cloning purposes. *Sporisorium reilianum* strains used in this study are listed in supplementary table 4. They are derivatives of the haploid solopathogenic strain JS161 which is capable of plant colonization without the need of a mating partner, because it expresses a compatible pheromone/receptor pair (Schirawski et al. 2010) *Escherichia coli* was grown in dYT liquid medium (1.6 % (w/v) Trypton, 1.0 % (w/v) Yeast Extract (Difco), 0.5 % (w/v) NaCl) or YT solid medium (0.8 % (w/v) Trypton, 0.5 % (w/v) Yeast-Extract, 0.5 % (w/v) NaCl, 1.3 % (w/v) agar) supplemented with 100 mg/mL Ampicillin when needed. The yeast *S. cerevisiae* was maintained in YPD solid medium (1 % (w/v) yeast extract, 2 % (w/v) Bacto-Pepton, 2 % (w/v) Bacto-Agar, 2 % (w/v) glucose) and grown on SC URA^-^ medium (1.7 % (w/v) Yeast Nitrogen Base without ammonium sulfate, 0.147 % (w/v) dropout-mix without Uracil, 2 % (w/v) glucose) for selecting transformants containing the plasmid pRS426 (Sikorski and Hieter 1989) (kindly provided by M. Bölker, Marburg) or derivatives of pRS426. Strains of *S. reilianum* were grown in liquid YEPS_light_ medium (1.0 % (w/v) yeast extract, 0.4% (w/v) peptone, 0.4% (w/v) sucrose) at 28°C on a rotary shaker at 200 rpm.

### Construction of S. reilianum strains

Polymerase chain reactions were performed using the Phusion High-Fidelity DNA Polymerase (New England Biolabs). Templates were either JS161 genomic or indicated plasmid DNA. Restriction enzymes were obtained from New England Biolabs. Protoplastmediated transformation was used to transform *S. reilianum* following a method established for *U. maydis* (*Schulz et al. 1990*). Transformants were selected on RegAgar plates (1.0 % (w/v) yeast extract, 0.4 % (w/v) Bacto-Pepton, 0.4 % (w/v) Sucrose, 1 M Sorbitol, 1.5 % (w/v) Bactoagar) supplemented with 200 μg/mL Geneticin and true resistance was tested by growing single colonies on PD plates (3.9 % (w/v) Potato-Dextrose Agar, 1 % (v/v) Tris-HCl (1M, pH 8.0)) supplemented with 50 μg/mL Geneticin. Gene replacements with resistance markers were generated with a PCR-based method employing the previously described *Sfi*I insertion cassette system (Brachmann et al. 2004; Kämper 2004) and were confirmed by Southern blot analysis. Genomic regions residing about 1 kb upstream (left border) or downstream (right border) adjacent to open reading frames of candidate genes were PCR-amplified using the listed primer pairs (supplementary table S5) and genomic DNA of JS161 as template. The resulting fragments were used for cloning plasmids containing the respective deletion constructs.

To obtain deletion constructs for the genes *sr10529* and *sr14347*, PCR fragments containing the left and right borders of each gene were ligated to the hygromycin resistance cassette of pBS-hhn (Kämper 2004) via *Sfi*I restriction sites and cloned into pCRII-TOPO (Life Technologies) to generate pTOPO Δsr10529 #1 and pTOPO Δsr14347 #1, respectively. Since the use of Geneticin as selection marker resulted in much less false positive transformants compared to the use of Hygromycin B, the hygromycin resistance cassettes in these plasmids were replaced by the Geneticin resistance cassette of pUMA 1057 (Brachmann et al. 2004) by ligation via *Sfi*I restriction sites, yielding plasmids pTOPO Δsr10529 G418 and pTOPO Δsr14347 Gen #1, respectively. Deletion constructs were PCR-amplified from plasmids pTOPO Δsr10529 G418 and pTOPO Δsr14347 Gen #1 using the listed primers (supplementary table S5) and used to transform the *S. reilianum* strain JS161 to generate the gene deletion strains JS161Δsr10529 and JS161Δsr14347, respectively.

The drag and drop cloning method in yeast (Jansen et al. 2005) was used to generate plasmids pRS426 Δsr12968 Hyg #1, pRS426 Δsr14944 Hyg #2, pRS426 Δsr10059 Hyg #1, pRS426 Δsr10182 Hyg #1, pRS426 Δsr14558 Hyg #1 and pRS426 Δsr12897 Hyg #1 which contain deletion constructs for deleting the candidate genes *sr12968*, *sr14944*, *sr10059*, *sr10182*, *sr14558* or *sr12897*. These plasmids are a derivate of plasmid pRS426, which can be maintained in *E. coli* and *S. cerevisiae* (Sikorski and Hieter 1989). PCR-amplified left and right borders of each candidate gene and the hygromycin resistance cassette were integrated in pRS426 by homologous recombination in *S. cerevisiae*. Subsequently, the hygromycin resistance cassette was replaced with the Geneticin resistance cassette by ligation via *Sfi*I restriction sites, yielding plasmids pRS426 Δsr12968 Gen #1, pRS426 Δsr14944 Gen #3, pRS426 Δsr10059 Gen #1, pRS426 Δsr10182 Gen #1, pRS426 Δsr14558 Gen #1 and pRS426 Δsr12897 Gen #5, respectively. Gene deletion constructs were PCR-amplified from the respective plasmid using listed primers (supplementary table S5). The obtained deletion constructs were used to transform the *S. reilianum* strain JS161 to generate the gene deletion strains JS161Δsr12968, JS161Δsr14944, JS161Δsr10059, JS161Δsr10182, JS161Δsr14558 and JS161Δsr12897, respectively.

The drag and drop cloning method was also used to generate plasmid pRS426 Δsr12084 Gen #1. PCR-amplified left and right borders of *sr12084* and the Geniticin resistance cassette were integrated in pRS426 by homologous recombination in *S. cerevisiae*. The gene deletion construct for deleting the candidate gene *sr12084* was PCR-amplified from plasmid pRS426 Δsr12084 Gen #1 using primers sr12084_lb_fw/sr12084_rb_rv and transformed into the *S. reilianum* strain JS161 to generate the gene deletion strain JS161Δsr12084.

### Virulence assays

The solopathogenic strain JS161 and derivatives thereof were grown in YEPS_light_ liquid medium to an optical density at 600 nm (OD_600_) of 0.8 - 1.0 and cell cultures were adjusted to an OD_600_ of 1.0 with sterile water prior to injection into one-week old maize (*Zea mays*) seedlings of the dwarf cultivar ‘Gaspe Flint’ (kindly provided by B. Burr, Brookhaven National Laboratories and maintained by self-pollination). Plants were sowed in T-type soil of ‘Fruhstorfer Pikiererde’ (HAWITA, Vechta, Germany) and grown in a temperature-controlled greenhouse (14h-/10h-light/dark cycle, with 28/20°C and 25,000 – 90,000 lux during the light period). Virulence symptoms were scored nine to ten weeks post infection according to previously described symptoms (Ghareeb et al. 2011) and the following categories were distinguished: the plant did not develop ears, the plant developed healthy ears shorter or equal to 1 cm or the plant developed healthy ears longer than 1 cm, the plant developed spiky ears, phyllody in ears or phyllody in tassels. Spore formation was only observed occasionally, and rarely the plant died due to the infection. Three independent infections were carried out per strain, mock treated plants were infected with water as control and at least three independent deletion strains were tested for virulence.

## Results

### Candidate effector genes are less conserved between species compared to other genes

We reconstructed families of homologous genes for the five smut fungi *U. hordei*, *U. maydis*, *S. scitamineum*, *S. reilianum* f. sp. *zeae* and *S. reilianum* f. sp. *reilianum* using the SiLiX clustering algorithm (Miele et al. 2011). We optimized the clustering parameters to maximize the occurrence of orthologues and minimize the number of paralogues within each family. In this way, we were able to reconstruct 8,761 families, among which 5,266 had at least one gene in each species (supplementary table S6). As a consequence, we found at least one homologous sequence in four species for 78 % of all genes. 5,254 gene families are found to have exactly one member in each species and were therefore taken as true orthologues (referred to as “core orthologous set” in the following). Considering that secreted proteins are putative effectors, we used SignalP (Petersen et al. 2011) to predict secretion of the encoded protein for each gene (supplementary table S1). We report that 920 (11 %) families contained only genes encoding a predicted secreted protein, while 7,657 (87 %) contained only genes encoding a protein not predicted to be secreted. The remaining 184 (2 %) families contained both predicted secreted and cytoplasmic proteins (supplementary table S6). The occurrence of families with both secreted and cytoplasmic proteins can be explained by (1) false negative predictions for secretion, as truncated C-terminal sequences were not removed from the data set, (2) wrong gene annotations or (3) gain or loss of a secretion signal peptide during effector evolution (Poppe et al. 2015). Among all predicted secreted proteins, 52 % have at least one orthologue in all other species, which is significantly less than the global 78 % proportion for all proteins (Chi-squared test, *P* value < 2.2×10^−16^). Genes encoding putative effector proteins are therefore less conserved across species than other genes, either because their sequence is evolving faster, preventing the recovery of homologous relationships, or because effector genes are created or lost at a higher rate. In *U. hordei*, we observe several species-specific family expansions. There were 17 families which encompassed five to 25 members, but no orthologue in other species (supplementary table S6). Moreover, we identified three families with up to 62 members in *U. hordei*, but only one member in up to three of the other species (supplementary table S6). Gene duplications in *U. hordei* have been hypothesized to be driven by mobile elements (Laurie et al. 2012).

### The genomes of S. reilianum f. sp. zeae and S. reilianum f. sp. reilianum diverged around one million years ago

To establish a frame for our comparative analysis we first calculated sequence similarity of the five smut fungi for non-coding intergenic and protein sequences. In addition, we estimated divergence times by performing a molecular dating analysis based on the core orthologues set of the five pathogens, using advanced models of protein sequence evolution and Bayesian inference as implemented in the PhyloBayes package (Lartillot et al. 2009). As calibration point, we used the divergence time of *U. hordei* and *U. maydis*, previously estimated to be between 27 and 21 Myr (Bakkeren and Kronstad, 2007). In alignable intergenic regions *U. hordei* shares 57 % identity with *S. reilianum* f. sp. *reilianum* and 77 % identity in protein sequences (fig. 1B). Monte-Carlo Markov chains were run for three independent gene samples totaling more than 20,000 amino acid positions each, and two chains were run in each case to assess convergence. The resulting posterior distribution of divergence times were used to infer 95 % posterior intervals. The split between *U. maydis* and the *Sporisorium* species was estimated to have occurred around 20 Myr ago (95 % posterior interval 25 to 12 Myr; fig 1A and supplementary table S2). *Sporisorium reilianum* f. sp. *reilianum* shares 61 % nucleotide identity in alignable intergenic regions with *U. maydis*, and 79 % sequence identity at the protein level (fig. 1B). The divergence times of *S. scitamineum* and the two formae speciales of *S. reilianum* were calculated to be 13 Myr ago (95 % posterior interval 19 to 7 Myr; fig. 1A and supplementary table S2), which is consistent with the mean divergence estimated between the hosts sorghum and sugarcane (10 Myr with a posterior interval of 8 to 13 Myr, average over eight studies, source: timetree.org (Kumar et al. 2017)). *Sporisorium reilianum* f. sp. *reilianum* and *S. scitamineum* share 74 % non-coding nucleotide identity and 88 % identity at the protein level (fig. 1B). Finally, the two *S. reilianum* strains diverged 1.1 Myr ago (95 % posterior interval 2.4 to 0.4 Myr; fig. 1A) and share 98 % noncoding nucleotide identity and 99 % protein identity (fig. 1B). We note that the estimation of this divergence date varied with the gene set used, and was in some cases found to be older (1.7 Myr, with a 95 % posterior interval of 4.7 to 0.6 Myr, see supplementary table S2). The comparison of the five smut genomes therefore encompasses a broad evolutionary time, and the divergence times obtained are compatible with previous estimates from smaller data sets (Munkacsi et al., 2007). The speciation times of the investigated smut species largely predate the 10,000 years of crop plant domestication, which implies that adaptation to the agricultural host, if any, will be negligible when interpreting the inter-specific patterns of sequence divergence, as it represents a marginal proportion of the time since the divergence from the ancestral species.

### Sporisorium reilianum contains the largest number of positively selected genes

To detect positive selection, 6,205 families with at least three members (orthologues and/or paralogues, see supplementary table S6) were, regardless of their species composition, aligned on the codon and amino acid level and a phylogenetic tree was inferred. Obtaining accurate alignments is critical for detecting positive selection since alignment errors frequently inflate the false discovery rate (Schneider et al. 2009; Jordan and Goldman 2012). We therefore developed a stringent bioinformatics pipeline for the filtering of sequence alignments by masking ambiguous alignment positions for further analysis (see Methods). To scan for positive selection, we employed a non-homogeneous model of sequence evolution allowing d_N_/d_S_ ratios to vary along the phylogeny, in combination with two heuristic model selection procedures to avoid over-parametrization issues (Nielsen and Yang 1998; Dutheil et al. 2012). Model parameters could not be fitted by either one of the two methods in 1.7% of branches. The two model selection procedures led to highly consistent estimates of branch-specific d_N_/d_S_ ratios (Spearman’s rank correlation coefficient equal to 0.85, p-value < 2.2×10^−16^). The distribution of d_N_ / d_S_ was highly skewed with a median value of 0.06, demonstrating the strong predominance of purifying selection throughout lineages and genes. The mean value of d_N_ / d_S_ ratios for lineages undergoing positive selection (d_N_ / d_S_ >1) was 4.1 (median 1.9). While a d_N_ / d_S_ ratio above one is indicative of positive selection, the absolute value of the ratio is a poor indicator of the strength of undergoing selection. In particular, high ratio values can be obtained because of low d_S_ values, and the d_N_ rate might include neutral substitutions, such as non-synonymous substitution that are conservative regarding certain biochemical properties of the amino-acids involved (Sainudiin et al. 2005). The largest number of genes with signs of positive selection was found in *S. reilianum* f. sp. *zeae* (84 genes, of which 25 encode predicted secreted proteins) and *S. reilianum* f. sp. *reilianum* (111 genes of which 27 encode predicted secreted proteins) (fig. 1C). In addition, a substantial number of positively selected candidate genes was also found in *U. hordei* (49, and of these, 22 genes are predicted to code for secreted proteins), but only very few in *U. maydis* (2 genes) and *S. scitamineum* (7 genes) (fig. 1C). A list of all proteins with their associated d_N_/d_S_ ratios in each species is provided in supplementary table S3. Predicted secreted proteins were significantly enriched in the group of proteins under positive selection in *U. hordei* and in the two investigated formae speciales of *S. reilianum* (*P* values < 10^−5^; Fisher’s exact test). This corroborates results of earlier studies in other pathosystems that showed that predicted secreted proteins are often under positive selection, which can be attributed to their direct interaction with host proteins (Joly et al., 2010; Wicker et al. 2013; Poppe et al., 2015). Notably, all genes found under positive selection in the two strains of *S. reilianum*, in *S. scitamineum* and in *U. maydis* share orthologous genes in the other species (supplementary tables S3 and S6). In contrast, genes with signs of positive selection in *U. hordei* belong largely (36 out of 49 genes) to families showing species-specific expansions (supplementary tables S3 and S6).

### Genes under positive selection in U. hordei are associated with uncharacterized interspersed repeats

Among the species compared here, the genome of *U. hordei* shows the highest fraction of repetitive elements (Laurie et al. 2012; Dutheil et al. 2016). Such elements are known to contribute to gene family expansions (Kazazian 2004), and have been suggested to contribute to adaptation by providing advantageous mutations, for instance by repeat-induced point mutations (RIP) leakage which was revealed in a species complex of *Leptosphaeria* (Rouxel et al. 2011; Grandaubert et al., 2014). As sequence signatures of RIP were found in LTR elements of *U. hordei* (Laurie et al., 2012), we tested whether genes under positive selection in *U. hordei* are physically associated with repetitive elements. We performed a binary logistic regression with the prediction of positive selection as a response variable (that is, whether the underlying branch has a d_N_/d_S_ ratio higher than one) and we considered three putative explanatory variables for each analyzed gene: (1) whether the gene is predicted to encode a secreted protein, (2) whether the gene is duplicated and (3) the distance of the gene to the closest interspersed repeat. The complete linear model explains 50 % of the observed variance, and the three explanatory variables are all significant at the 0.1 % level (supplementary table S7). These results suggest that positively selected genes in *U. hordei* are associated with duplication events, and positive selection is more likely to occur at genes encoding putative effectors. In addition, the proximity of interspersed repeats increases the odds of positive selection, independently of the two other effects, and is confirmed by a stratification approach: the effect still holds when only duplicated genes are considered, or only genes encoding a secreted protein, or the combination of the two (supplementary table S7). This finding corroborates previous results obtained in other microbial plant pathogens where it was described that effector genes tend to localize in repeat rich regions and where it was suggested that such regions contribute to the rapid evolution of effector genes (Raffaele and Kamoun, 2012).

### Positively selected genes encoding cytoplasmic proteins in S. reilianum and U. hordei

While we expect effector genes to be under positive selection, we find that the majority of positively selected genes in *S. reilianum* encodes cytoplasmic proteins (fig. 1C). To assess the putative functional role of these genes, we performed a Gene Ontology term enrichment analysis, comparing cytoplasmic proteins under positive selection to cytoplasmic proteins not under positive selection (table 1). This analysis revealed that genes with a potential role in metabolic processes, like sulfur compound metabolism, molybdopterin cofactor metabolic process, RNA metabolic process, organic cyclic compound metabolic process and oxidoreductase activity, as well as responses to starvation and extracellular stimuli are significantly overrepresented at the 5% level (Fisher’s classic test with Parent-Child correction; see table 1). This could indicate that cytoplasmic proteins under positive selection contribute to metabolic changes which might be needed to survive with the limited nutrients available on the surface or in the biotrophic interface of different host plants. A similar analysis for cytoplasmic proteins under positive selection in *U. hordei* was conducted, but only led to top-level categories (DNA integration, DNA metabolic process and isomerase activity; see table 1).

**Table 1:** Gene Ontology terms significantly overrepresented in positively selected genes encoding cytoplasmic proteins in *S. reilianum* and *U. hordei*

| Gene Ontology Id | Gene Ontology Description | Category<sup>a)</sup> | P values<sup>b)</sup> | Species<sup>c)</sup> |
| --- | --- | --- | --- | --- |
| GO:0030532 | small nuclear ribonucleoprotein complex | CC | 0.023 | Srr |
| GO:0045263 | proton-transporting ATP synthase complex, coupling factor F(o) | CC | 0.038 and 0.036 | Srr and Srz |
| GO:0044425 | membrane part | CC | 0.047 | Srr |
| GO:0031668 | cellular response to extracellular stimulus | BP | 0.015 and 0.013 | Srr and Srz |
| GO:0051186 | cofactor metabolic process | BP | 0.015 | Srr |
| GO:0042594 | response to starvation | BP | 0.017 and 0.012 | Srr and Srz |
| GO:0009267 | cellular response to starvation | BP | 0.020 and 0.014 | Srr and Srz |
| GO:0006790 | sulfur compound metabolic process | BP | 0.022 | Srr |
| GO:0051301 | cell division | BP | 0.023 | Srr |
| GO:0006777 | Mo-molybdopterin cofactor biosynthetic process | BP | 0.027 | Srr |
| GO:0009605 | response to external stimulus | BP | 0.027 and 0.012 | Srr and Srz |
| GO:0043545 | molybdopterin cofactor metabolic process | BP | 0.034 | Srr |
| GO:0043413 | macromolecule glycosylation | BP | 0.036 | Srr |
| GO:0022402 | cell cycle process | BP | 0.037 | Srr |
| GO:0051189 | prosthetic group metabolic process | BP | 0.039 | Srr |
| GO:0006139 | nucleobase-containing compound metabolic process | BP | 0.012 | Srz |
| GO:0016070 | RNA metabolic process | BP | 0.023 | Srz |
| GO:1901360 | organic cyclic compound metabolic process | BP | 0.016 | Srz |
| GO:0006725 | cellular aromatic compound metabolic process | BP | 0.024 | Srz |
| GO:0046483 | heterocycle metabolic process | BP | 0.023 | Srz |
| GO:0035383 | thioester metabolic process | BP | 0.041 | Srr |
| GO:1902589 | single-organism organelle organization | BP | 0.048 | Srr |
| GO:0015074 | DNA integration | BP | 0.039 | Uh |
| GO:0006259 | DNA metabolic process | BP | 0.023 | Uh |
| GO:0010181 | FMN binding | MF | 0.047 and 0.022 | Srr and Srz |
| GO:0030515 | snoRNA binding | MF | 0.021 | Srz |
| GO:0016491 | oxidoreductase activity | MF | 0.028 | Srz |
| GO:0016853 | isomerase activity | MF | 0.029 | Uh |

<sup>a)</sup> CC, Cellular Component; BP, Biological Process; MF, Molecular Function.
<sup>b)</sup> calculated by Fisher's classic test with Parent-Child correction. Only entries with $P$ value ≤ 0.05 are shown.
<sup>c)</sup> Species in which GO terms were found enriched. Srr, *S. reilianum* f. sp. *reilianum*; Srz, *S.* *reilinaum* f. sp. *zeae*; Uh, *U. hordei*.

### Virulence contribution of effector genes showing signs of positive selection in S. reilianum

Candidate effector genes inferred to be under positive selection in a particular species could play a critical role in pathogenicity. Therefore, we sought to assess the contribution to virulence of such candidate genes by creating individual deletion mutants. In total, we tested nine candidate genes with high d_N_/d_S_ ratios and predicted to encode secreted proteins: three with signatures of positive selection only in *S. reilianum* f. sp. *zeae*, three with signatures of positive selection in *S. reilianum* f. sp. *zeae* as well as in *S. reilianum* f. sp. *reilianum* and three with signatures of positive selection only in *S. reilianum* f. sp. *reilianum*. All nine chosen candidate genes together with their characteristics are summarized in table 2. Deletion mutants were generated in the haploid solopathogenic strain JS161 of *S. reilianum* f. sp. *zeae*. This strain is capable of colonizing maize plants and cause disease without a compatible mating partner (Schirawski et al. 2010) but virulence is much reduced relative to infection with mating-compatible wild-type strains and spores are only rarely produced. Deletion mutants were also generated in strain JS161 in cases where positive selection was only detected in *S. reilianum* f. sp. *reilianum* (table 2), because no solopathogenic strain is presently available for *S. reilianum* f. sp. *reilianum*. For each gene at least three independent deletion mutants were generated and tested for virulence. To determine virulence, Gaspe Flint, a dwarf variety of corn, was infected and symptoms were scored in male and female flowers (fig. 2). Only the deletion of *sr10529*, a gene showing positive selection in both formae speciales of *S. reilianum*, showed a strong reduction in virulence (table 2 and fig. 2). The gene *sr10529* in *S. reilianum* f. sp. *zeae* is orthologous to the previously identified and characterized gene *pit2*(*UMAG_01375*) in *U. maydis*.

**FIG. 2.**
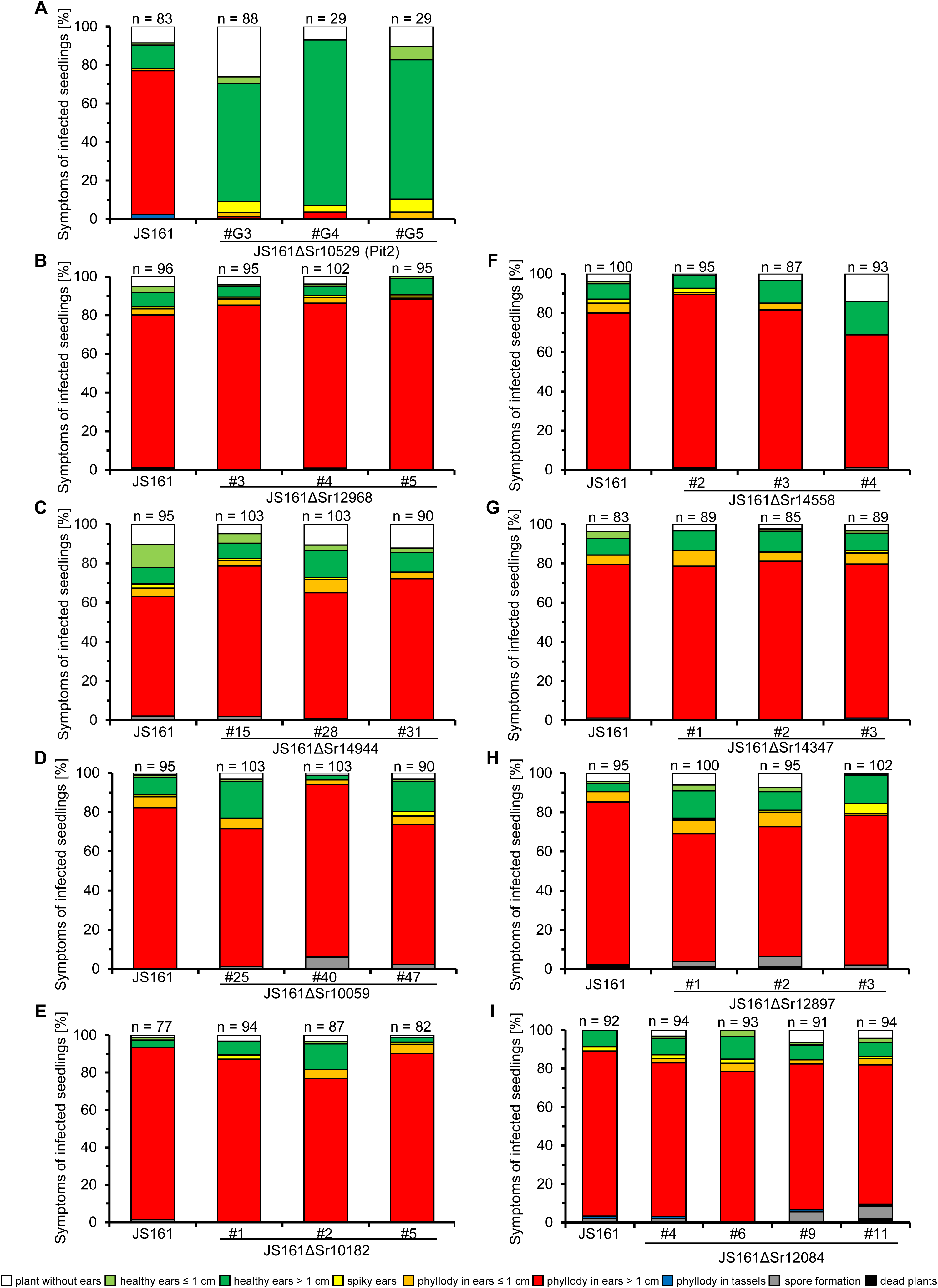
Virulence phenotypes of single deletion mutants of positively selected genes in *S. reilianum* f. sp. *zeae*. Plants of the maize variety ‘Gaspe Flint’ were infected with the solopathogenic strain JS161 or independent deletion mutants of candidate genes as indicated below each bar. Deletion of *sr10529* led to a strong reduction in virulence (A). In contrast, deletion of the candidate genes *sr12968* (B), *sr14944* (C), *sr10059* (D), *sr10182* (E), *sr14558* (F), *sr14347* (G), *sr12897*(H), and *sr12084* (I) did not alter virulence. Symptoms were scored about 9 weeks post infection and categorized according to severeness as illustrated in the legend below the bar plots. Results are shown as mean of three independent experiments in relation to the total number of infected plants, which is indicated above each bar (n). Note that strains JS161ΔSr10529 #G4 and #G5 (A) were only infected in one replicate.

**Table 2:** Positively selected genes that were deleted in *S. reilianum* f. sp. z*eae* and 984 their selection criteria

| Family | Gene Id | Gene description | d<sub>N</sub>/d<sub>S</sub> | Species with detected positive selection<sup>a)</sup> | Number of Paralogs in Srz<sup>b)</sup> | Virulence phenotype of candidate gene deletion | Closest ortholog in *U. maydis*<sup>c)</sup> |
| --- | --- | --- | --- | --- | --- | --- | --- |
| FAM001428 | *sr10529* | conserved hypothetical protein | 31.1469 | Srz and Srr | 0 | virulence abolished | *UMAG_01375* (*pit2*)<sup>6)</sup> |
| FAM005472 | *sr10059* | conserved hypothetical *Ustilaginaceae*-specific protein | 6.53881 | Srz and Srr | 0 | virulence unaffected | *UMAG_05306* (cluster 19A)<sup>7)</sup> |
| FAM000532 | *sr10182* | conserved hypothetical protein | 1.57473 | Srr | 10<sup>3)</sup> | virulence unaffected | *UMAG_00492* |
| FAM002067 | *sr12968* | conserved hypothetical protein | 37.9007 | Srr | 0 | virulence unaffected | *UMAG_02006* |
| FAM003728 | *sr14558* | conserved hypothetical protein | 24.355 | Srz | 0 | virulence unaffected | *UMAG_03564* |
| FAM004113 | *sr14944* | conserved hypothetical *Ustilaginaceae*-specific protein | 4.30527 | Srz and Srr | 0 | virulence unaffected | *UMAG_04034* (cluster 11-16)<sup>8)</sup> |
| FAM003465 | *sr14347* | conserved hypothetical protein | 544.37 | Srz | 5<sup>4)</sup> | virulence unaffected | *UMAG_03349* |
| FAM001868 | *sr12897* | conserved hypothetical protein | infinite<sup>1)</sup> | Srr | 0 | virulence unaffected | *UMAG_01820* |
| FAM000842 | *sr12084* | conserved hypothetical protein | infinite<sup>2)</sup> | Srz | 2<sup>5)</sup> | virulence unaffected | *UMAG_00792* (cluster 1-32)<sup>8)</sup> |

<sup>a)</sup> species are *S. reilianum* f. sp. *zeae* (Srz) and *S. reilianum* f. sp. *reilianum* (Srr) <sup>b)</sup> based on blastp search with an e-value cutoff of 0.001 <sup>c)</sup> based on blastp search <sup>1)</sup> infinity due to low value of $d_s$ <sup>2)</sup> infinity due to long branch for *sr12085*, a species-specific duplicate in *S. reilianum* f. sp. *zeae* <sup>3)</sup> the ten paralogs include: *sr13431*, *sr11876*, *sr16607*, *sr11405*, *sr10621*, *sr16723*, *sr16877*, *sr13293*, *sr11163.2* and *sr15970* <sup>4)</sup> the five paralogs include: *sr12257*, *sr11661*, *sr13976*, *sr14607* and *sr11273* <sup>5)</sup> the two paralogs include: *sr12085* and *sr12086* <sup>6)</sup> as described in Doehlemann et al., 2011 and Mueller et al., 2013 <sup>7)</sup> as described in Kämper et al., 2006 <sup>8)</sup> as described in Schirawski et al., 2010

Pit2 plays an essential role in virulence as inhibitor of a group of maize papain-like cysteine proteases that are secreted to the apoplast (Doehlemann et al. 2011; Mueller et al. 2013).Previous work identified a conserved domain of 14 amino acids (PID14) in Pit2 as required and sufficient for the inhibition of maize cysteine proteases (Mueller et al. 2013). When the branch-site model of PAML 4 (Yang 2007) was used to identify amino acid residues under positive selection in the Pit2 orthologues of the two *S. reilianum* species, only two residues residing in the PID14 domain were found under positive selection. However, 24 positively selected residues were detected outside this domain in the 57 amino acid long C-terminal part (fig. 3).

**FIG. 3.**
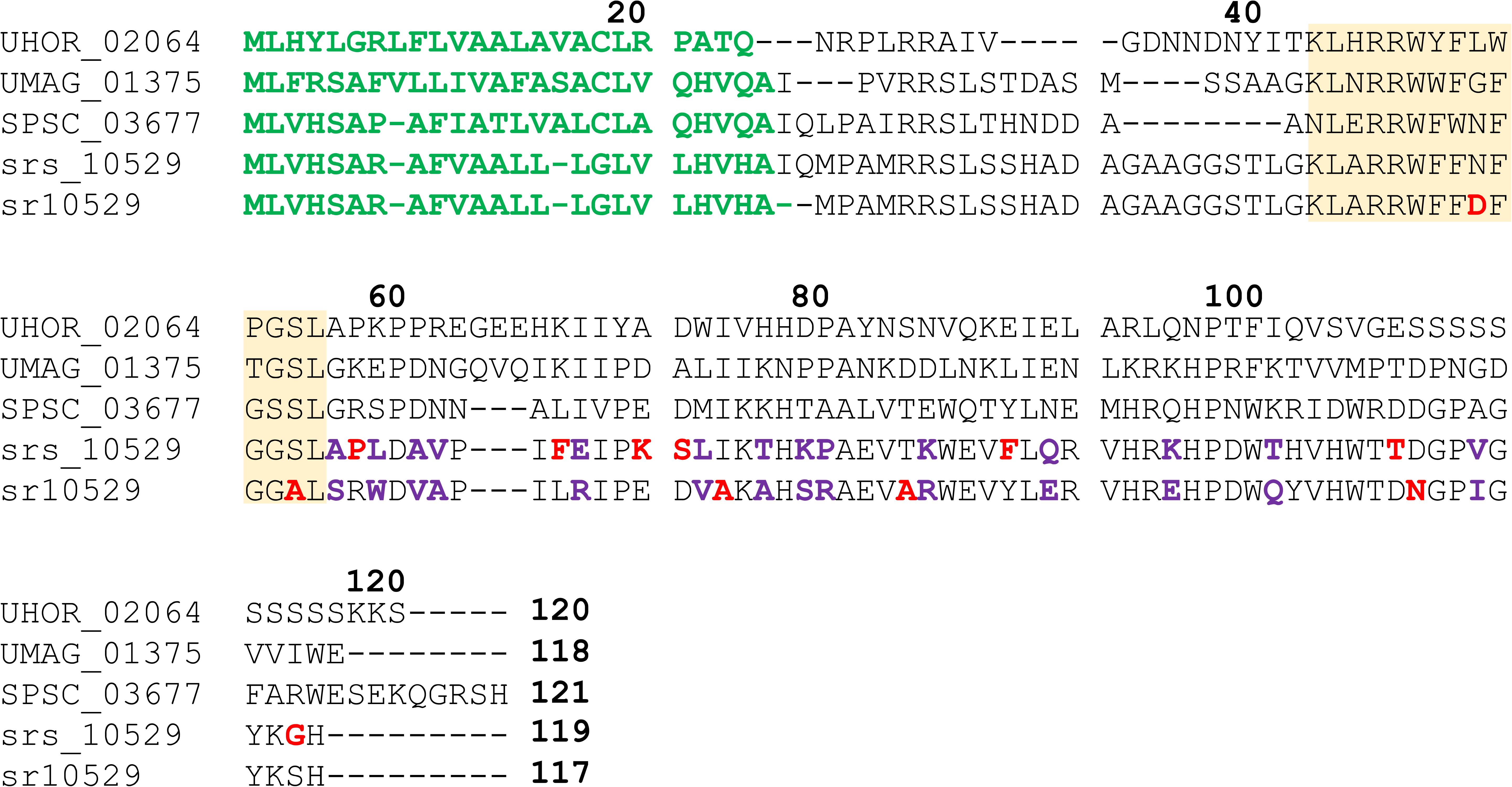
Distribution of positively selected amino acids in the cysteine protease inhibitor Pit2. The alignment shows the protein sequences of orthologues in *U. hordei* (UHOR_02064), *U. maydis* (UMAG_01375), *S. scitamineum* (SPSC_03677), *S. reilianum* f. sp. *zeae* (sr10529) and *S. reilianum* f. sp. *reilianum* (srs_10529). Sites under positive selection detected by a branch-site model are indicated by colored bold letters. Residues colored in red indicate positive selection detected in the respective species and purple residues indicate sites found under positive selection in both species. The yellow shaded area is orthologous to the previously identified conserved PID14 domain, which is required and sufficient for inhibition of a group of papain-like cysteine proteases. Green sequences indicate secretion signal peptides and bold numbers above the alignment indicate positions in UHOR_02064.

## Discussion

We used evolutionary comparative genomics of five related smut fungi infecting four different host plants to identify genes with a signature of positive selection during species divergence, with a focus on genes encoding predicted secreted proteins, as such genes were suggested to contribute to virulence in various plant pathogenic microbes (Aguileta et al. 2010; Stukenbrock et al. 2011; Hacquard et al. 2012; Dong et al. 2014; Huang et al. 2014; Sharma et al. 2014) Our analysis revealed that positive selection is found between paralogous genes in *U. hordei*, where they belong to families with species-specific expansions. In contrast, genes under positive selection in the other four species belong to families of orthologous sequences. While we find evidence for a large set of genes under positive selection in the *S. reilianum* species, signatures for positive selection are hardly detectable in the more distant relatives *U. hordei*, *U. maydis* and *S. scitamineum* that diverged earlier. Finding evidence for positive selection over time spans of several millions of years is notoriously difficult (Gillespie 1994) because of two main reasons: (1) periods where genes are evolving under positive selection may occur episodically and may be followed by long episodes of purifying selection, leading to an average d_N_/d_S_ below 1 on long periods of time and (2) fast evolving genes may diverge to an extent where their homology is difficult to infer and where they can no longer be aligned reliably. To overcome this problem, more genome information of species with intermediate branching points is needed (Gillespie 1994).

Predicted secreted proteins were about three times overrepresented in the set of positively selected genes, which illustrates the importance of secreted proteins in adaptation processes of smut fungi. This also corroborates results in other plant pathogenic microbes like *Melampsora sp., Z. tritici* and the wheat powdery mildew *Blumeria graminis* (Joly et al. 2010; Stukenbrock et al. 2011; Wicker et al. 2013). However, the majority of positively selected genes encodes cytoplasmic proteins (fig. 1C), suggesting that both secreted and non-secreted proteins are important targets of adaptation. A Gene Ontology analysis in *S. reilianum* showed that mainly processes related to metabolism and its regulation as well as responses to starvation and external stimuli are enriched in cytoplasmic proteins under positive selection. This points at a role of these proteins in adaptation to differences in nutrient availability in the respective host plants maize and sorghum as well as responses to cues originating from the respective host (Haueisen and Stukenbrock 2016). A study conducted in *U. maydis* has shown that the fungus induces major metabolic changes in the host plant upon infection during establishment of biotrophy and undergoes a series of developmental transitions during host colonization that are likely influenced by the host environment (Doehlemann et al. 2008). It is thus conceivable that the two *S. reilianum* accessions have adapted to their different hosts that differ significantly for example in their amino acid and vitamin composition (Etuk et al. 2012). Furthermore, recent studies in *U. maydis* suggested that intracellular changes of metabolism influence virulence, and therefore the underlying proteins could be targets of positive selection (Kretschmer et al. 2012; Goulet et al. 2017).

Out of nine deletions of positively selected genes, only one mutant, lacking *sr10529*, was affected in virulence. While six of the deleted genes are single genes in *S. reilianum* f. sp. *zeae* for which we failed to identify paralogs, *sr12084* has two paralogs, *sr14347* has five paralogs and *sr10182* has ten paralogs. We restricted our analyses to generating deletion mutants in some of the genes under positive selection. This leaves open the possibility that the paralogous genes have redundant functions in virulence. Adapting the CRISPR-Cas9 technology allowing multiplexing (Schuster et al. 2017) to *S. reilianum* will be instrumental in testing this hypothesis in future studies. Alternatively, the candidate effectors we investigated may be needed under conditions which differ from those tested here. For example, *S. reilianum* f. sp. *zeae* can also systemically colonize maize plants via root infection (Mazaheri-Naeini et al. 2015), a colonization route we have not assessed in our experimental setup. Moreover, we employed only one maize cultivar for infection assays. Results from other pathosystems suggest that virulence effects can strongly depend on the host and pathogen genotypes, in particular in the presence of *R* and *avr* genes (Petit-Houdenot and Fudal, 2017). No *avr-R* gene interaction was described so far in the *S. reilianum* f. sp. zeae-maize pathosystem. Instead, quantitative virulence differences are observed when different host cultivars are infected (Lübberstedt et al. 1999). Knowing the expression profile of effector genes may assist the identification of differences in development of the mutants compared to wild type strains. Since we lack this information, we scored disease symptoms only in the inflorescences about nine weeks after infection. Additionally, it may be possible that small differences in virulence between JS161 and deletion mutants of candidate genes remain undetected due to the weak infection behavior of JS161. For example, spore formation is only rarely observed after infecting maize plants with the solopathogenic strain. In contrast, infections resulting from infections with two compatible haploid strains show spores in about 40 % of the infected plants (Zuther et al. 2012). This means that defects related to spore formation will not be evident in mutants of JS161. In three cases positive selection was detected in orthologous genes in *S. reilianum* f. sp. *reilianum* while candidate effector genes were for experimental reasons deleted in *S. reilianum* f. sp. *zeae*. Therefore, it cannot be excluded that these effectors might have a virulence function in *S. reilianum* f. sp. *reilianum*. In this case, the positively selected effector genes might have evolved during adaptation to the sorghum host and present host specificity genes. In summary, our virulence assays leave open the possibility that the eight candidate genes which did not show a contribution to virulence could play a role in pathogenicity under conditions not tested here. Alternatively, candidate effector proteins might also be positively selected for traits that are not directly linked to pathogenicity. Such traits could for instance involve competition with large numbers of other plant colonizing microbes (Zhan and McDonald 2013; Rovenich et al. 2014). Secreted proteins of *S. reilianum* could act for example as toxin or could efficiently utilize resources from the environment and thereby limit the growth of other microbes. In these cases, a contribution to virulence is not expected to be observed in the employed infection assay with the effector gene mutants. Moreover, our molecular dating analysis showed that the common ancestors of the investigated smut species originated before the beginning of crop domestication. Therefore, positive selection, whose signs we detect by our approach, has most likely occurred on ancestral host plants and not on the domesticated host maize. Consequently, some of the candidate effector genes under positive selection might not be important for the colonization of crop plants, but for infection of related wild species.

In *U. maydis*, we note that effector genes residing in clusters whose deletion affected virulence (Kämper et al. 2006; Schirawski et al. 2010) have similar d_N_/d_S_ ratios as effector genes in clusters where the deletion had no effect on virulence (median d_N_/d_S_ ratio 0.0619 vs.0.1094; Wilcoxon rank test with *P* value = 0.1848). Furthermore, orthologues of the effectors Pep1, Stp1 and Cmu1, which were shown to have important roles in pathogenicity of *U. maydis* (Djamei et al. 2011; Doehlemann et al. 2009; Schipper 2009) showed no signature of positive selection. These observations could suggest that certain fungal effector proteins are under evolutionary constraint and are therefore not free to accumulate non-synonymous mutations. Such effectors are conserved over long time spans (Schirawski et al. 2010; Hemetsberger et al. 2015; Sharma et al. 2015) and this illustrates that they are instrumental for successful infections in a large group of smut fungi. They probably target molecules shared by several host plants, for example housekeeping functions that cannot easily evolve in response to the binding of an effector.

One candidate gene (*sr10529*) under positive selection in both formae speciales of *S. reilianum* showed a strong contribution to virulence upon deletion. It is orthologous to the previously described protease inhibitor Pit2 in *U. maydis*, where the deletion also abolished virulence (Doehlemann et al. 2011; Mueller et al. 2013). Positively selected residues in the PID14 domain of Pit2 might reflect that different proteases need to be inhibited in maize and sorghum. Pit2 might thus contribute to determining the host range of the respective species. A role of cysteine protease inhibitors in host specificity was demonstrated in *Phytophthora infestans*, a pathogen of potato and its sister species *Phytophthora mirabilis*, which infects the ornamental plant *Mirabilis jalapa*. Positively selected orthologous protease inhibitors were shown to inhibit proteases specific to the respective host plants and this specificity could be traced back to a single amino acid substitution (Dong et al. 2014). Surprisingly, 24 positively selected sites in Pit2 were detected outside the PID14 domain in the 57 amino acid long C-terminal part in both *S. reilianum* f. sp. *zeae* and *S. reilianum* f. sp. *reilianum*. This finding raises the intriguing possibility that the C-terminus of Pit2 might possess a second function that is independent of protease inhibition. Earlier work has shown that the *pit1* gene encoding a transmembrane protein is located next to the *pit2* effector gene and both genes contribute similarly to virulence (Doehlemann et al. 2011). Furthermore, *pit1* and *pit2* are divergently transcribed, which makes it likely that the expression of *pit1* and *pit2* is co-regulated. In addition, this gene arrangement of *pit1* and *pit2* is conserved in *U. hordei*, *U. maydis*, *S. scitamineum* and *S. reilianum* (Sharma et al. 2015). This finding has led to the speculation that Pit1 and Pit2 somehow act together to govern virulence of *U. maydis* and related smut fungi. It was hypothesized that that Pit2 shuttles apoplastic maize proteins towards Pit1, thereby scavenging damage-associated molecules (Doehlemann et al. 2011). In this scenario, the positively selected amino acids in the C-terminus of Pit2 could have been selected for scavenging such molecules as adaptation to the two hosts. In future studies it will be highly interesting to complement the *pit2* mutant of *S. reilianum* f. sp. *zeae* with the *pit2* orthologue of *S. reilianum* f. sp. *reilianum* to see if this promotes virulence on sorghum.

## Conclusions

Screens for genes with signs of positive selection are commonly used to identify candidate effector genes in various plant pathogenic microbes. However, it is currently largely open whether positively selected effector genes play indeed a role in virulence. Here, we used comparative genomics of five smut fungi and showed that only one out of nine genes under positive selection contributes to virulence of *S. reilianum*. Moreover, the majority of positively selected genes did not encode predicted secreted proteins. Our results leave open the possibility that many genes with signatures of positive selection contribute to virulence under conditions not tested in this study or are selected in traits that are not directly related to pathogenicity.

## Acknowledgements

Our work was supported through the LOEWE program of the state of Hesse through SYNMICRO and through the Max Planck Society. GS was funded by the International Max Planck Research School (IMPRS) for Molecular, Cellular and Environmental Microbiology. We thank Michael Bölker for providing the yeast strain BY4741 and the plasmid pRS426 which were used for cloning purposes, and Benoit Nabholz and Nicolas Lartillot for guidance with the dating analysis. We are grateful to Markus Rampp (Max Planck Computing Center Garching) for assistance with computational work and to Stefan Schmidt for expert greenhouse service, as well as all present and former group members for lively discussions and Eva H. Stukenbrock for her comments on the manuscript.

## List of supplementary tables

**Supplementary table S1:** Sources of genomic data for the five smut fungi species investigated

**Supplementary table S2:** Results of the molecular dating analysis

**Supplementary table S3:** Prediction of secretion, Gene Ontology Terms and d_N_ / d_S_ ratios for all 33,940 proteins in five smut fungi

**Supplementary table S4:** List of strains of *S. reilianum* f. sp. *zeae* used in the present study

**Supplementary table S5:** List of primers used in the present study

**Supplementary table S6:** Grouping of 33,940 proteins of five compared smut fungi species in 8,761 families

**Supplementary table S7:** Results of a linear model illustrating the associations between gene duplications, positive selection and candidate effector genes

## List of supplementary figures

**Supplementary figure 1.** Number of families consisting of 1:1 orthologues in relation to varying settings for coverage and identity in the clustering program SiLiX. The maximum number of families containing 1:1 orthologues can be obtained with a coverage between 5 % and 45 % and an identity of 40 %.

**Supplementary figure 2.** Trace of the Monte-Carlo Markov chains for 3 gene samples (see Methods). Vertical lines show the burning phase (10,000 iterations)

## References

Anikster Y. 1984. The Formae Speciales. In: Bushnell WR and Roelfs AP (Eds.) The cereal rusts. Academic Press (London).

Aguileta G, et al. 2012. Genes under positive selection in a model plant pathogenic fungus, *Botrytis*. Infect Genet Evol 12: 987–996.

Aguileta G, et al. 2010. Finding candidate genes under positive selection in non-model species: examples of genes involved in host specialization in pathogens. Mol Ecol 19: 292–306.

Aguileta G, Refregier G, Yockteng R, Fournier E, Giraud T. 2009. Rapidly evolving genes in pathogens: methods for detecting positive selection and examples among fungi, bacteria, viruses and protists. Infect Genet Evol 9: 656–670.

Alexa A, Rahnenführer J, Lengauer T. 2006. Improved scoring of functional groups from gene expression data by decorrelating GO graph structure. Bioinformatics 22: 1600–1607.

Altschul SF, Gish W, Miller W, Myers EW, Lipman DJ. 1990. Basic local alignment search tool. J Mol Biol 215: 403–410.

Bagchi R, et al. 2014. Pathogens and insect herbivores drive rainforest plant diversity and composition. Nature 506: 85–88.

Bakkeren G, Kronstad JW. 2007. Bipolar and Tetrapolar Mating Systems in the Ustilaginales. In: Heitman J, Kronstad JW, Taylor J, Casselton L (eds). Sex in fungi. ASM Press, Washington.

Begerow D, et al. 2014. Ustilaginomycotina. In: Esser K (ed). The Mycota - A comprehensive treatise on fungi as experimental systems for basic and applied research. Systematics and evolution (Part A). Springer, Heidelberg.

Blanchette M, et al. 2004. Aligning multiple genomic sequences with the Threaded Blockset Aligner. Genome Res 14: 708–715.

Brachmann A, König J, Julius C, Feldbrügge M. 2004. A reverse genetic approach for generating gene replacement mutants in *Ustilago maydis*. Mol Genet Genomics 272: 216–226.

Brown JKM, Tellier A. 2011. Plant-parasite coevolution: bridging the gap between genetics and ecology. Annu Rev Phytopathol 49: 345–367.

Charif D, Lobry JR. 2007. SeqinR 1.0-2: a contributed package to the R project for statistical computing devoted to biological sequences retrieval and analysis. In: Bastolla U, Porto M, Roman HE, Vendruscolo M, editors. Structural approaches to sequence evolution: Molecules, networks, populations. New York: Springer Verlag. p. 207–232.

Dean R, et al. 2012. The top 10 fungal pathogens in molecular plant pathology. Mol Plant Pathol 13: 414–430.

de Jonge R, Bolton MD, Thomma BP. 2011. How filamentous pathogens co-opt plants: the ins and outs of fungal effectors. Curr Opin Plant Biol 14: 400–406.

Djamei A, et al. 2011. Metabolic priming by a secreted fungal effector. Nature 478: 395–398.

Doehlemann G, Reissmann S, Aßmann D, Fleckenstein M, Kahmann R. 2011. Two linked genes encoding a secreted effector and a membrane protein are essential for *Ustilago maydis-*induced tumour formation. Mol Microbiol 81: 751–766.

Doehlemann G, et al. 2009. Pep1, a secreted effector protein of *Ustilago maydis*, is required for successful invasion of plant cells. PLoS Pathog 5: e1000290.

Doehlemann G, et al. 2008. Reprogramming a maize plant: transcriptional and metabolic changes induced by the fungal biotroph *Ustilago maydis*. Plant J 56: 181–195.

Dong S, et al. 2014. Effector specialization in a lineage of the Irish potato famine pathogen. Science 343: 552–555.

Dutheil JY, Boussau B. 2008. Non-homogeneous models of sequence evolution in the Bio++ suite of libraries and programs. BMC Evol Biol 8: 255.

Dutheil JY, et al. 2012. Efficient selection of branch-specific models of sequence evolution. Mol Biol Evol 29: 1861–1874.

Dutheil JY, et al. 2016. A tale of genome compartmentalization: the evolution of virulence clusters in smut fungi. Genome Biol Evol 8: 681–704.

Etuk EB, et al. 2012. Nutrient composition and feeding value of *Sorghum* for livestock and poultry: a review. J Anim Sci Adv 2: 510–524.

Fisher MC, et al. 2012. Emerging fungal threats to animal, plant and ecosystem health. Nature 484: 186–194.

Franceschetti M, et al. 2017. Effectors of filamentous plant pathogens: commonalities amid diversity. Microbiol Mol Biol R 81: e00066-00016.

Gehrig H, Schussler A, Kluge M. 1996. *Geosiphon pyriforme*, a fungus forming endocytobiosis with *Nostoc* (cyanobacteria), is an ancestral member of the *Glomales:* evidence by SSU rRNA analysis. J Mol Evol 43: 71–81.

Ghareeb HA, Becker A, Iven T, Feussner I, Schirawski J. 2011. *Sporisorium reilianum* infection changes inflorescence and branching architectures of maize. Plant Physiol 156: 2037–2052.

Giraldo MC, Valent B. 2013. Filamentous plant pathogen effectors in action. Nat Rev Microbiol 11: 800–814.

Gillespie JH. 1994. The causes of molecular evolution. Oxford: Oxford University Press.

Goulet KM, Saville BJ. 2017. Carbon acquisition and metabolic changes during fungal biotrophic plant pathogenesis: insights from Ustilago maydis. Can J Plant Pathol 39: 247–266.

Grandaubert J, et al. 2014. Transposable element-assisted evolution and adaptation to host plant within the Leptosphaeria maculans-Leptosphaeria biglobosa species complex of fungal pathogens. BMC Genomics 15: 891.

Grossmann S, Bauer S, Robinson PN, Vingron M. 2007. Improved detection of overrepresentation of Gene-Ontology annotations with parent child analysis. Bioinformatics 23: 3024–3031.

Guindon S, et al. 2010. New algorithms and methods to estimate maximum-likelihood phylogenies: assessing the performance of PhyML 3.0. Syst Biol 59: 307–321.

Hacquard S et al. 2012. A Comprehensive Analysis of Genes Encoding Small Secreted Proteins Identifies Candidate Effectors in *Melampsora larici-populina* (Poplar Leaf Rust). Mol Plant Microbe In 25: 279–293.

Harrell Jr. FE. 2015. Regression Modeling Strategies. Springer Series in Statistics, Switzerland.

Haueisen J, Stukenbrock EH. 2016. Life cycle specialization of filamentous pathogens - colonization and reproduction in plant tissues. Curr Opin Microbiol 32: 31–37.

Hemetsberger C, et al. 2015. The fungal core effector Pep1 is conserved across smuts of dicots and monocots. New Phytol 206: 1116–1126.

Huang J, Si W, Deng Q, Li P, Yang S. 2014. Rapid evolution of avirulence genes in rice blast fungus *Magnaporthe oryzae*. BMC Genetics 15: 45.

Jansen G, Wu C, Schade B, Thomas DY, Whiteway M. 2005. Drag&Drop cloning in yeast. Gene 344: 43–51.

Joly DL, Feau N, Tanguay P, Hamelin RC. 2010. Comparative analysis of secreted protein evolution using expressed sequence tags from four poplar leaf rusts (Melampsora spp.). BMC Genomics 11: 422.

Jordan G, Goldman N. 2012. The effects of alignment error and alignment filtering on the sitewise detection of positive selection. Mol Biol Evol 29: 1125–1139.

Kämper J. 2004. A PCR-based system for highly efficient generation of gene replacement mutants in *Ustilago maydis*. Mol Genet Genomics 271: 103–110.

Kämper J, et al. 2006. Insights from the genome of the biotrophic fungal plant pathogen *Ustilago maydis*. Nature 444: 97–101.

Kazazian HH. 2004. Mobile elements: drivers of genome evolution. Science 303: 1626–1632.

Khrunyk Y, Münch K, Schipper K, Lupas AN, Kahmann R. 2010. The use of FLP-mediated recombination for the functional analysis of an effector gene family in the biotrophic smut fungus *Ustilago maydis*. New Phytol 187: 957–968.

Kretschmer M, Klose J, Kronstad JW. 2012. Defects in mitochondrial and peroxisomal β- oxidation influcence virulence in the maize pathogen Ustilago maydis. Eukaryot Cell 11: 1055–1066.

Kumar S, Stecher G, Suleski M, Hedges SB. 2017. TimeTree: a resource for timelines, timetrees, and divergence times. Mol Biol Evol 34: 1812–1819.

Lanver D, et al. 2017. *Ustilago maydis* effectors and their impact on virulence. Nat Rev Microbiol 15: 409–421.

Lartillot N, Lepage T, Blanquart S. 2009. PhyloBayes 3: a Bayesian software package for phylogenetic reconstruction and molecular dating. Bioinformatics 25: 2286–2288.

Laurie JD, et al. 2012. Genome comparison of barley and maize smut fungi reveals targeted loss of RNA silencing components and species-specific presence of transposable elements. Plant Cell 24: 1733–1745.

Le SQ, Gascuel O. 2008. An improved general amino acid replacement matrix. Mol Biol Evol 25: 1307–1320.

Lo Presti L, et al. 2015. Fungal effectors and plant susceptibility. Annu Rev Plant Biol 66: 513–545.

Löytynoja A, Goldman N. 2008. Phylogeny-aware gap placement prevents errors in sequence alignment and evolutionary analysis. Science 320: 1632–1635.

Lübberstedt T, Xia XC, Tan G, Liu X, Melchinger AE. 1999. QTL mapping of resistance to *Sporisorium reiliana* in maize. Theor Appl Genet 99: 593–598.

Martin FM, Uroz S, Barker DG. 2017. Ancestral alliances: Plant mutualistic symbioses with fungi and bacteria. Science 356: eaad4501.

Martinez-Espinoza AD, Garcia-Pedrajas MD, Gold SE. 2002. The Ustilaginales as plant pests and model systems. Fungal Genet Biol 35: 1–20.

Mazaheri-Naeini M , Sabbagh SK, Martinez Y, Séjalon-Delmas N, Roux C. 2015. Assessment of *Ustilago maydis* as a fungal model for root infection studies. Fungal Biol 119: 145–153.

Miele V, Penel S, Duret L. 2011. Ultra-fast sequence clustering from similarity networks with SiLiX. BMC Bioinformatics 12: 116.

Möller M, Stukenbrock EH. 2017. Evolution and genome architecture in fungal plant pathogens. Nat Rev Microbiol 15: 756–771.

Mueller AN, Ziemann S, Treitschke S, Aßmann D, Doehlemann G. 2013. Compatibility in the *Ustilago maydis-*maize interaction requires inhibition of host cysteine proteases by the fungal effector Pit2. PLoS Pathog 9: e1003177.

Munkacsi AB, Stoxen S, May G. 2007. Domestication of maize, sorghum, and sugarcane did not drive the divergence of their smut pathogens. Evolution 61: 388–403

Nielsen R. 2005. Molecular signatures of natural selection. Annu Rev Genet 39: 197–218.

Nielsen R, Yang Z. 1998. Likelihood models for detecting positively selected amino acid sites and applications to the HIV-1 envelope gene. Genetics 148: 929–936.

Parniske M. 2008. Arbuscular mycorrhiza: the mother of plant root endosymbioses. Nat Rev Microbiol 6: 763–775.

Petersen TN, Brunak S, von Heijne G, Nielsen H. 2011. SignalP 4.0: discriminating signal peptides from transmembrane regions. Nat Methods 8: 785–786.

Petit-Houdenot Y, Fudal I. (2017). Complex Interactions between Fungal Avirulence Genes and Their Corresponding Plant Resistance Genes and Consequences for Disease Resistance Management. Front Plant Sci 8: 1072.

Plissonneau C, et al. 2017. Using population and comparative genomics to understand the genetic basis of effector-driven fungal pathogen evolution. Front Plant Sci 8: 119.

Poppe S, Dorsheimer L, Happel P, Stukenbrock EH. 2015. Rapidly evolving genes are key players in host specialization and virulence of the fungal wheat pathogen *Zymoseptoria tritici (Mycosphaerella graminicola)*. PLoS Pathog 11: e1005055.

Que Y, et al. 2014. Genome sequencing of *Sporisorium scitamineum* provides insights into the pathogenic mechanisms of sugarcane smut. BMC Genomics 15: 996.

Raffaele S, Kamoun S. 2012. Genome evolution in filamentous plant pathogens: why bigger can be better. Nat Rev Microbiol 10: 417–430.

Ranwez V, Harispe S, Delsuc F, Douzery EJP. 2011. MACSE: Multiple Alignment of Coding SEquences accounting for frameshifts and stop codons. PLOS One 6: e22594.

Remy W, Taylor TN, Hass H, Kerp H. 1994. Four hundred-million-year-old vesicular arbuscular mycorrhizae. PNAS 91: 11841–11843.

Romiguier J, et al. 2012. Fast and robust characterization of time-heterogeneous sequence evolutionary processes using substitution mapping. PLOS One 7: e33852.

Rouxel T, et al. 2011. Effector diversification within compartments of the *Leptosphaeria maculans* genome affected by Repeat-Induced Point mutations. Nat Commun 2: 202.

Rovenich H, Boshoven JC, Thomma BPHJ. 2014. Filamentous pathogen effector functions: of pathogens, hosts and microbiomes. Curr Opin Plant Biol 20: 96–103.

Sainudiin R, et al. 2005. Detecting site-specific physicochemical selective pressures: applications to the Class I HLA of the human major histocompatibility complex and the SRK of the plant sporophytic self-incompatibility system. J Mol Evol 60: 315–326.

Schipper K 2009. Charakterisierung eines *Ustilago maydis* Genclusters, das für drei neuartige sekretierte Effektoren kodiert. Philipps-Universität Marburg.

Schirawski J, et al. 2010. Pathogenicity determinants in smut fungi revealed by genome comparison. Science 330: 1546–1548.

Schneider A, et al. 2009. Estimates of positive Darwinian selection are inflated by errors in sequencing, annotation, and alignment. Genome Biol Evol 1: 114–118.

Schulz B, et al. 1990. The *b* alleles of *U. maydis*, whose combinations program pathogenic development, code for polypeptides containing a homeodomain-related motif. Cell 60: 295–306.

Schuster M, Schweizer G, Kahmann R. 2017. Comparative analyses of secreted proteins in plant pathogenic smut fungi and related basidiomycetes. Fungal Genet Biol in press.

Schuster M, Schweizer G, Reissmann S, Kahmann R. 2016. Genome editing in *Ustilago maydis* using the CRISPR-Cas system. Fungal Genet Biol 89: 3–9.

Sharma R, Mishra B, Runge F, Thines M. 2014. Gene loss rather than gene gain is associated with a host jump from monocots to dicots in the smut fungus *Melanopsichium pennsylvanicum*. Genome Biol Evol 6: 2034–2049.

Sharma R, Xia X, Riess K, Bauer R, Thines M. 2015. Comparative genomics including the early-diverging smut fungus *Ceraceosorus bombacis* reveals signatures of parallel evolution within plant and animal pathogens of fungi and oomycetes. Genome Biol Evol 7: 2781–2798.

Sikorski RS, Hieter P. 1989. A system of shuttle vectors and yeast host strains designed for efficient manipulation of DNA in *Saccharomyces cerevisiae*. Genetics 122: 19–27.

Sperschneider J, et al. 2015. Genome-wide analysis in three *Fusarium* pathogens identifies rapidly evolving chromosomes and genes associated with pathogenicity. Genome Biol Evol 7: 1613–1627.

Sperschneider J, et al. 2014. Diversifying selection in the wheat stem rust fungus acts predominantly on pathogen-associated gene families and reveals candidate effectors. Front Plant Sci 5: 372.

Stergiopoulos I, de Wit PJ. 2009. Fungal effector proteins. Annu Rev Phytopathol 47: 233–263.

Stukenbrock EH, et al. 2011. The making of a new pathogen: insights from comparative population genomics of the domesticated wheat pathogen *Mycosphaerella graminicola* and its wild sister species Genome Res 21: 2157–2166.

Taniguti LM, et al. 2015. Complete genome sequence of *Sporisorium scitamineum* and biotrophic interaction transcriptome with sugarcane. PLOS One 10: e0129318.

Tellier A, Moreno-Gámez S, Stephan W. 2014. Speed of adaptation and genomic footprints of host-parasite coevolution under arms race and trench warefare dynamics. Evolution 68: 2211–2224.

Thompson JD, Plewniak F, Poch O. 1999. A comprehensive comparison of multiple sequence alignment programs. Nucleic Acids Res 27: 2682–2690.

Thorne JL, Kishino H, Painter IS. 1998. Estimating the rate of evolution of the rate of molecular evolution. Mol Biol Evol 15: 1647–1657.

Tiffin P, Moeller DA. 2006. Molecular evolution of plant immune system genes. Trends Genet 22: 662–670.

Toruño TY, Stergiopoulos I, Coaker G. 2016. Plant-pathogen effectors: cellular probes interfering with plant defenses in spatial and temporal manners. Annu Rev Phytopathol 54: 419–441.

Wicker T, et al. 2013. The wheat powdery mildew genome shows the unique evolution of an obligate biotroph. Nat Genet 45: 1092–1096.

Yang Z. 1998. Likelihood ratio tests for detecting positive selection and application to primate lysozyme evolution. Mol Biol Evol 15: 568–573.

Yang Z, Nielsen R. 1998. Synonymous and nonsynonymous rate variation in nuclear genes of mammals. J Mol Evol 46:409–418.

Yang Z. 2006. Computational Molecular Evolution. Oxford Series in Ecology and Evolution, Oxford University Press.

Yang Z. 2007. PAML 4: phylogenetic analysis by maximum likelihood. Mol Biol Evol 24: 1586–1591.

Zhan J, McDonald BA. 2013. Experimental measures of pathogen competition and relative fitness. Annu Rev Phytopathol 51: 131–153.

Zuther K, et al. 2012. Host specificity of *Sporisorium reilianum* is tightly linked to generation of the phytoalexin luteolinidin by *Sorghum bicolor*. Mol Plant Microbe Interact 25: 1230–1237.

